# Museum insights for conservation: Unraveling the Extinction Factors in the Jambato Harlequin Frog, *Atelopus ignescens*

**DOI:** 10.1101/2024.12.07.627320

**Authors:** Mónica A. Guerra, Chengchen Gao, Samuel Crickenberger, Michelle Vélez, Luis A. Coloma, Silu Wang

**Author notes:** Corresponding authors: Mónica A. Guerra,; Silu Wang.

## Abstract

Natural history museums harbor invaluable resources for conserving endangered species by providing insights into the mechanism of historical population declines. Here we conducted data synthesis to better understand the extinction factors of the iconic Jambato Harlequin frog, *Atelopus ignescens*, which was widespread in the Ecuadorian Andes before 1985 but vanished in 1988. The mechanism of extinction remains elusive, as it was uncertain whether *Batrachochytrium dendrobatidis (Bd)* fungus infection, climate change, or the interaction of the two factors primarily contributed to the rapid population decline. Here we synthesize historical data from natural history museums, the Global Biodiversity Information Facility (GBIF), and mitochondrial DNA sequences to uncover the factors that have potentially contributed to the extinction of the Jambato Harlequin frog. We found excessive rare alleles reflected in the negative Tajima’s D estimated from the mitochondrial DNA samples collected in 1984, which indicates a selective sweep or bottleneck in the population shortly before the historical extinction. There was a marginal effect of the time before versus during *Bd* epizootics on the body sizes of the adult males, which was relatively weaker than the sex effect and geographic effect. The body size of adult males, but not the females form a geographic cline where the individuals in the northeast were larger than the males in the southwest. Species distribution modeling based on temperature and precipitation accurately predicted the occupancy of *A. ignescens* in 1960-69 (R^2^ = 0.72, accuracy = 0.9). This model further predicted the rapid decline in species distribution over decades of climate change between 1970-2020, which could have contributed to the rapid population decline in the 1980s. Collectively, our data synthesis revealed a strong climate effect and a weak epizootics effect on *A. ignescens* extinction, which unravels the mysterious rapid population decline and informs ongoing conservation efforts. This investigation inspires future museum genomic studies to dissect the potential climatic maladaptation behind one of the most dramatic extinction events in modern history.

## Introduction

Amidst the catastrophic loss of global vertebrate biodiversity with an average of 69% decline in wild populations since 1970 (Living Planet Index, WWF 2022), 40.7% of amphibian species are threatened (Luedke et al. 2023). Multiple factors drive amphibian declines, and their significance varies among species, populations, and regions (Campbell-Grant et al. 2016, 2020; Luedke et al 2023). Specifically, the decline in Andean anuran populations has been predominantly attributed to the interplay between chytridiomycosis, a disease caused by the fungus *Batrachochytrium dendrobatidis (Bd)*, and climate change (Pounds et al. 2006; Cohen et al. 2019). The susceptibility of Andean anurans to *Bd* fungus infection seems to be heightened due to their habitat maintaining the optimal growth temperature range (17-25 °C) for *Bd* fungus (Piotrowski et al. 2004; Stevenson et al. 2013; Woodhams et al. 2008).

The Jambato Harlequin frog, *Atelopus ignescens*, is an anuran species that was once widespread in the Ecuadorian Andes, with reports of 10 to 50 individuals per square meter in the early 1980s (Black 1982; Valle 1984; Almendáriz and Orcés 2004; Coloma and Duellman in press). However, its population sharply declined in the late 1980s, and the last individual was seen in 1988 (Coloma et al 2000). Despite extensive surveys from 1999 to 2003 yielding no results, a relict population was rediscovered in Northern Ecuador in 2016 (Coloma 2016). Since then, this population formed the basis of an ex-situ breeding program aimed at conserving the potentially last remaining individuals of the species. A field survey conducted in 2016 and 2019 confirmed the presence of some individuals within this population infected with *Bd*. Even with the presence of *Bd*, this population remains the only known wild population of *A. ignescens* (Jaynes et al. 2022; Vega-Yánez 2024).

Museum data provides organismal resources for conservation, offering invaluable insights into species distributions, functional morphological evolution, and historical population trends. In this context, we investigate the Jambato Harlequin Frog, *A. ignescens*, as a case study to elucidate the abundance of data attainable from museum specimens, highlighting the importance of museum collections in conservation biology. Such specimens are particularly significant in the face of threats like climate change and pathogens, which have been proposed as the main causes of precipitous population declines and extinctions in several species of Andean anurans (Ron et al. 2003; Merino-Viteri et al. 2005; Menéndez-Guerrero and Graham 2013). Museum specimens collected over the decades before the extinction contain valuable information on sex ratios, age structure, morphology, pathogen genetics, and population genetics, which empowers the dissection of the mechanisms underlying historical population changes.

Here we used curated museum and GBIF data to examine the possible influence of *Bd* fungus, annual temperature, and precipitation, on the decline of *A. ignescens* during the 1980s. To explore the potential factors behind the rapid population decline, we constructed a species distribution model, examined the patterns of rare alleles with mitochondrial DNA, and measured adult body size from museum specimens. If these climate variables are significant predictors of *A. ignescens* distribution, we predict a dramatic decline in the species’ predicted range following the climate changes that occurred after 1970. Previous research indicates that smaller frogs are more susceptible to chytridiomycosis due to osmoregulatory and metabolic disruptions (Wu et al. 2018). This susceptibility may have led to directional selection for larger body size in *A. ignescens* during the *Bd* epizootics of the 1980s. If chytridiomycosis was a factor contributing to the species’ extinction, we would expect a significant increase in the average body size of *A. ignescens* during *Bd* epizootics in the 1980s. Although *Bd* is not directly evaluated in this study, body size data offers a comparative approach to examine morphological changes before and during the population decline, acknowledging that multiple factors may have contributed to the species’ decline. Additionally, we discuss how additional genomic studies could elucidate the additional knowledge gaps for the restoration and conservation of this iconic frog species.

## Methods

### Climate data

We acquired historical climate data from six decades: 1960-69, 1970-79, 1980-89, 1990-99, 2000-2010, and 2010-19 from WorldClim (Fick and Hijmans 2017; Harris et al. 2020). Specifically, we downloaded historical monthly climate data from CRU-TS-4.06 by using WorldClim 2.1 for bias correction with a spatial resolution of 2.5 minutes (∼21 km2 at the equator). The climate variables available at this temporal and spatial resolution were: (1) monthly minimum temperature (°C), (2) monthly maximum temperature (°C), and (3) total precipitation (mm). These variables were chosen to predict the distribution of *A. ignescens* based on previous studies modeling the effects of climate change on the distribution of other tropical frogs and factors known to strongly influence the physiology of other closely related species (Schivo et al. 2019).

### Museum specimens

To examine whether there was a change in body size before and during the *Bd* epizootics, we measured the Snout-Vent Length (SVL) and examined the sex of 176 adult museum specimens of *A. ignescens*. These individuals were collected from localities between 78.89°W and 78.08°W longitude, and 0.15°N to 1.51°S latitude, including 83 from the Herpetology Collection of the Natural History Museum Gustavo Orcés V. at Escuela Politécnica Nacional (MEPN-H), 87 from Museum of Pontificia Universidad Católica del Ecuador (QCAZ), and 6 from Kansas University Natural History Museum. In total, there were 45 females and 17 males before 1980, and 71 females and 43 males between 1980-1990. We considered males >29 mm SVL and females >30.5 mm as adults. We dissected most of the specimens to determine the sex based on gonad morphology. Few of them were not dissected because of apparent sexual landmarks such as visible eggs through the skin in females or the presence of secondary sexual traits, such as keratinized nuptial pads, vocal slits, or hypertrophic limbs in males. Based on field notes and personal communications of researchers from those periods we know most of them were collected by Visual Encounter Surveys (VES), although the sampling collecting methods for *A. ignescens* vary amongst collectors and projects and may include sampling plots or linear transects. Before the 1980s, all specimens in the collection were preserved in formalin 10%. During the first years of the 80s, wet specimens were transferred to ethanol 70%, and after the 80s specimens were fixed with formalin 10% and preserved in ethanol 70% (Almendáriz et al. 2023).

### Regression Model of Body Size

We examined spatial and temporal variation of body size in adult females and males. To effectively understand the spatial distribution of frog SVL, we calculated the one-dimensional Geographic Index as the Euclidian distance

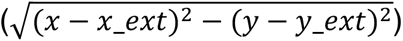 of each sampling location (x, y) to the sampling site at the southwest extreme (x_ext, y_ext). The southwest extreme site was regarded as the reference starting point so that this geographic index (GI) reflects the relative locations of *A. ignescens* along the Andes from southwest (low GI) to northeast (high GI) (Fig. 1). To test if there is spatial and/or temporal effect(s) on SVL in males and females, we conducted linear model selection. A full model was constructed with the response variable being SVL and the predicting variables being: (1) Geographic Index, (2), periods (before or after 1980), and (3) sex (female vs male), as well as all their interactions (i.e. lm (SVL ∼ geographic index * periods * sex). With this full model, we conducted backward stepwise model selection until the model with the lowest Akaike Information Criteria (AIC) was achieved (Hastie & Pregibon, 1992). The best model with the lowest AIC is further investigated. We calculated the partial effect sizes of the predictors in the selected model with the *rstatix* package (Kassambara, 2023) in R 4.3.2 (R Core Team, 2023). To test if the effect sizes of the predictors were significantly different from zero, we used bootstrapping. Briefly, we sampled the full dataset with replacement, fitted the selected regression model (above), and computed the effect sizes for all the predicting variables as the bootstrap distributions of the predictors. We evaluate whether the 95% confidence intervals of the bootstrap distributions were greater than zero. If so, the corresponding predictor has a significant effect.

**Fig. 1.**
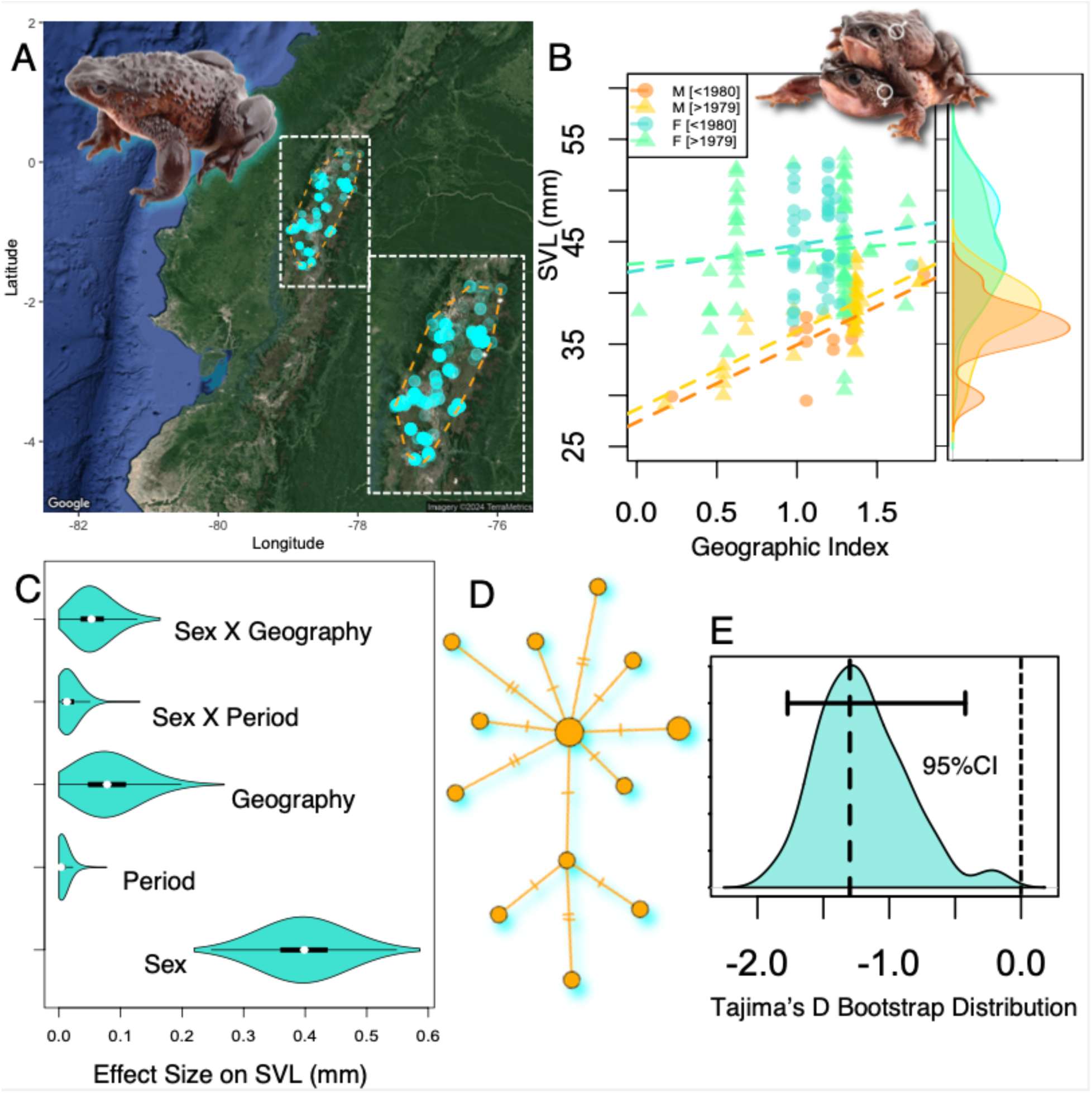
The population biology of *Atelopus ignescens* before extinction. **A**, Locations of species sighting before 1990 (N = 1122). **B-C**, The stepwise multiple regression model selection (Table S2) concluded with the selected model (**B**) for explaining the variation of SVL (mm) with the following predictors (**C**): (1) sex, (2) geography, (3) time, (4) interaction of sex and geography, and (5) the interaction of sex and time. **B**, adult females (F) were larger than the males (M). The density distributions of SVL of each sex before or during 1980 were shown on the right side. **D**, The haplotype network of mitochondrial DNA samples collected in 1984. Each circle refers to a haplotype, and each tick mark corresponds to a mutation in addition to the line connecting the haplotype circles. **E**, Bootstrap distribution of Tajima’s D was significantly negative, suggesting excessive rare alleles in the 1984 sample shortly before *A. ignescens* extinction in 1988.

### Genetic diversity prior to extinction

To understand the genetic diversity within *A. ignescens* around the onset of rapid population decline, we downloaded mitochondrial sequences from GenBank, where the mitochondrial NADH dehydrogenase subunit 2 gene and cytochrome oxidase subunit I gene were partially sequenced and tRNA-Trp, tRNA-Ala, tRNA-Asx, tRNA-Cys, tRNA-Tyr genes were completely sequenced for 16 individuals sampled in 1984 (Guayasamin et al. 2010). We downloaded the concatenated mitochondrial gene sequences from GenBank (with accession numbers listed in Table S1) for each individual and aligned the sequences with MUSCLE in MEGA11 (Tamura et al. 2021). We calculated the haplotype network with POPART (Leigh and Bryant 2015) with this dataset. To further evaluate the evolutionary forces shaping *A. ignescens* prior to the historical rapid population decline, we calculated the bootstrap distribution of Tajima’s D with the *pegas* package (Paradis 2010) of R. To compute the bootstrap distribution of Tajima’s D, we sampled the 1984 mtDNA dataset (Table S1) with replacement for 1000 iterations. We calculated Tajima’s D for each of the bootstrap samples to generate a bootstrap distribution of Tajima’s D. If the population was under the mutation-drift balance, the 95% confidence interval (CI) of Tajima’s D bootstrap distribution should not deviate from zero (Tajima, 1989). If the population was experiencing selective sweep or bottleneck, there should have been excessive rare alleles, thus the 95% CI of the Tajima’s D bootstrap distribution should be less than zero. Alternatively, if there was a deficit of rare alleles, the Tajima’s D 95% CI should be positive.

### Species Distribution Modeling

To test whether there were significant reductions in the distribution of *A. ignescens* in response to the change of temperature and precipitation between 1970-1990, we fit a GLM to the climate and *A. ignescens* data from 1960 to 1970. We assume that the *A. ignescens* population was in stable condition in the 1960-69 due to the stable natural history records of this species prior to the global climate shift in the 1970s (Black 1982; Valle 1984; Almendáriz and Orcés 2004; Sarkar & Maity, 2021). The *A. ignescens* presence data was acquired from the Global Biodiversity Information Facility (GBIF.org, Sep. 28th, 2023; https://doi.org/10.15468/dl.9uzx5v). We further included additional precise coordinate records of the museum samples listed above (Fig. 1 A). While GBIF provides a vast amount of data for researchers, there are several potential issues associated with it (bias in sampling efforts across regions and time, misidentifications, sampling errors leading to incomplete coverage). To correct for misidentification, we removed points outside the altitudinal range of 2483 to 4385 meters and longitudes greater than −60, as well as the points with latitudes more than 0.15 North and 1.51 South (Ron et al. 2003, and updates by Guayasamin et al. 2010, and Vega-Yánez et al. 2024). This resulted in N =1122 presence points (Fig. 1 A). To understand the distribution of *A. ignescens* when the population was stable, we further filtered to only include the presence points sampled between 1960 and 1970 (N=265). For the absence data, we randomly sampled 500 pseudo-absence points in the density trough (area with n<1, Fig. S2) of the *A. ignescens* density kernel within the square of - 80° to -77° W and -3° to 1° N, fitted as bivariate normal distribution with all the presence data of *A. ignescens* (Fig. 1 A, N=1122). We randomly divided the presence/absence data into 5 folds with ⅘ of the data being training data and the remainder ⅕ being the testing data to be used for model fitting and accuracy evaluation respectively (detailed below).

With the training data being the response variable, we fit a binomial generalized linear model (GLM) with the three climate variables measured in 1960-69 and their interactions being the individual predicting variables. To find the set of necessary predicting variables that effectively explain the response variable, we conducted backward stepwise model selection and concluded with the model with the lowest AIC (Table S3). In the selected model, the effect sizes of the regression coefficients were calculated with *eta_squared* function in R. With the testing data, we calculated the threshold probability of presence with *pa_evaluate* function in the *predicted* package in R (Fielding and Bell 1997; Liu et al. 2011), and prediction accuracy as (true positive + true negative)/total testing size. In addition, we evaluated the prediction accuracy of the 1960-69 model for predicting species presence in 1970-79 and 1980-89. To generate the testing data in each period, we combined the pseudo-absence data (see above) with the presence data from the corresponding decade and randomly selected ⅕ of the combined dataset as the testing dataset to calculate prediction accuracy.

To understand whether climate change affected the distribution of *A. ignescens* over time, we projected the 1960-69 GLM model (see above) onto climate measurements from each of the subsequent decades (1970-79, 1980-89, 1990-99, 2000-09, and 2010-19). We further tested whether there was a reduction of predicted presence with bootstrap sampling with 1000 iterations. Briefly, in each iteration, we sampled all the available sites with replacement and found the total number of sites with predicted *A. ignescens* presence over the total number of sites, as the *proportion of suitable habitats*. We computed the *proportion of suitable habitats* simultaneously for the predicted presence of each decade. Then we compared the 95% CIs of the *proportion of suitable habitats* bootstrap distribution across decades.

## Results

There was limited signature that Bd infection contributed to the population change in *A. ignescens*. The multiple regression model with time and the interaction of sex and time fitted the data better than the models without these predictors after controlling for the likelihood elevation with additional parameters (Table S2). This corresponds to increased intercepts in the SVL-geography regression line during 1980s Bd epizootics than before the 1980s in males but not in females (Table S3, Fig. 1 B). However, the effect size of time-related predictors was small: sex and time interaction had a mean effect size of 0.0167 (95% CI: 0.0002 to 0.0538), while the independent time effect size was only 0.0067 (95% CI: 7 × 10^-6^ to 0.0383) (Fig. 1 C).

In contrast, we discovered a relatively larger geographic effect size on SVL. The independent geographic effect size was 0.0818 (95% CI: 0.0112 to 0.1806) (Fig. 1 C). There was also a geography-sex interaction with an effect size of 0.0571 (95%CI: 0.0127 to 0.1319) (Fig. 1 B-C, S2). In particular, the SVL of males exhibited strong geographic clines in both periods, but the female SVL was not significantly correlated with geographic indices (Fig. 1 B, S2).

There were significantly excessive rare alleles in the 1984 mitochondrial genetic samples (Fig. 1 D-E) with the 95% confidence interval of the Tajima’D bootstrap distribution ranging less than 0. The haplotype network (Fig. 1 D) did not reflect population substructure (Fig. S3). This significantly negative Tajima’s D is consistent with bottleneck/selective sweep, which might be related to Bd infection and/or climate change in the early 1980s.

The climate variables effectively predicted the distribution of *A. ignescens* in 1960-69 (R^2^ = 0.72, Table 1). Testing data confirmed the 90% accuracy of this GLM model. The accuracy for predicting species distribution in 1970-79 and 1980-89 declined to 78% and 62%, respectively. The minimum temperature (effect size = 0.57, 95% CI: 0.53-1), precipitation (effect size = 0.15, 95% CI: 0-1), and the interaction of minimum and maximum temperatures (effect size = 0.15, 95% CI: 0.11-1) were key predictors for *A. ignescens* distribution. We further predicted the *A. ignescens* presence with this 1960-69 model and the climate data collected from the subsequent decades.

**Table 1.**
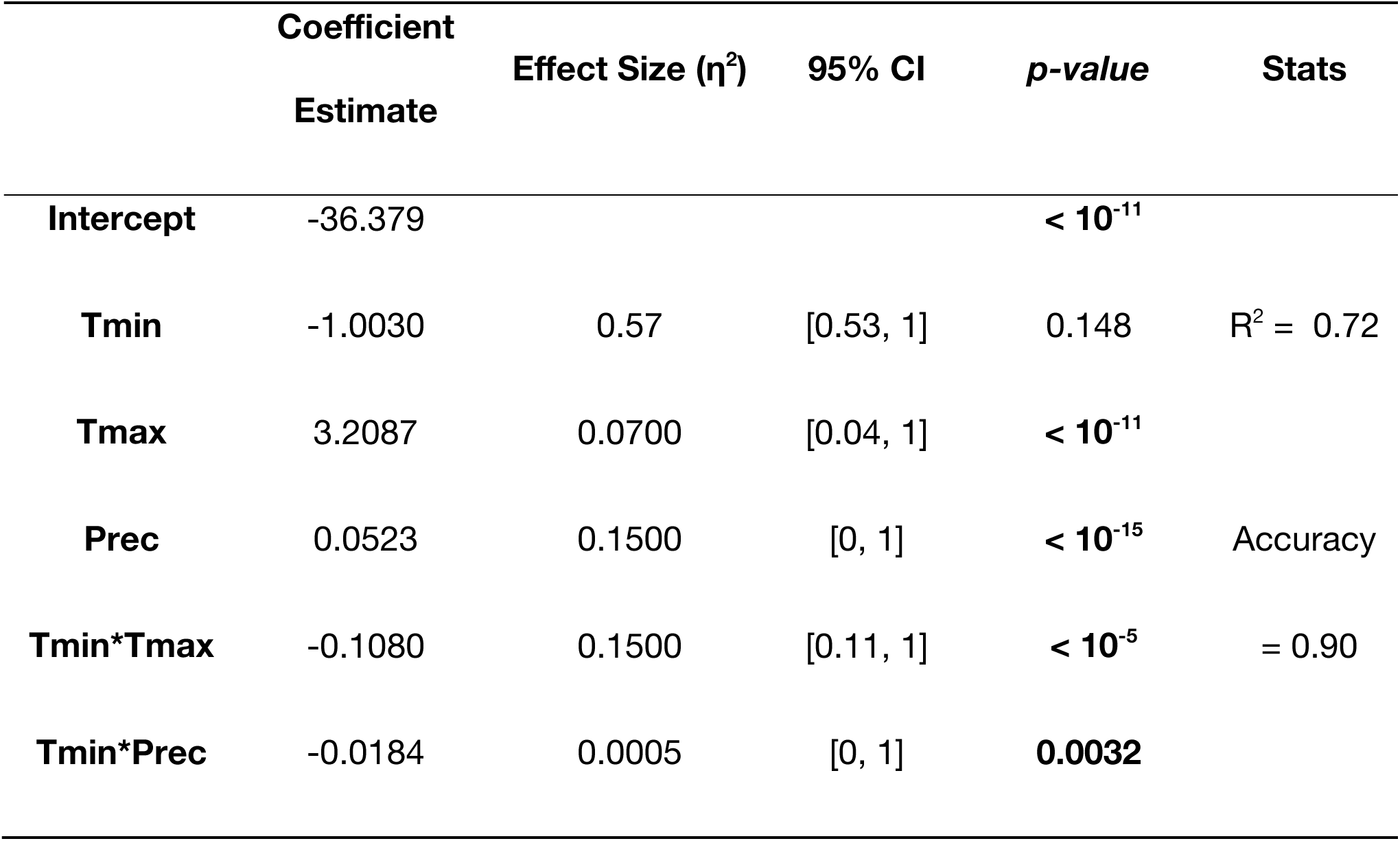
Regression coefficients of the species distribution model for 1960-69. Stepwise model selection resulted in the model with minimum (T_min_), maximum temperatures (T_max_), precipitation (Prec), and the interaction of T_min_ and T_max_, as well as the interaction of T_min_ and Prec.

There were significant (*p* < 0.001, repeated measures ANOVA, Table S4) changes in temperature (Fig. 2) and precipitation (Fig. 3) in the Ecuadorian Andes in 1970-2020, which resulted in a drastic shrinkage of predicted presence in 1970-79 and onward (Fig. 4).

**Fig. 2.**
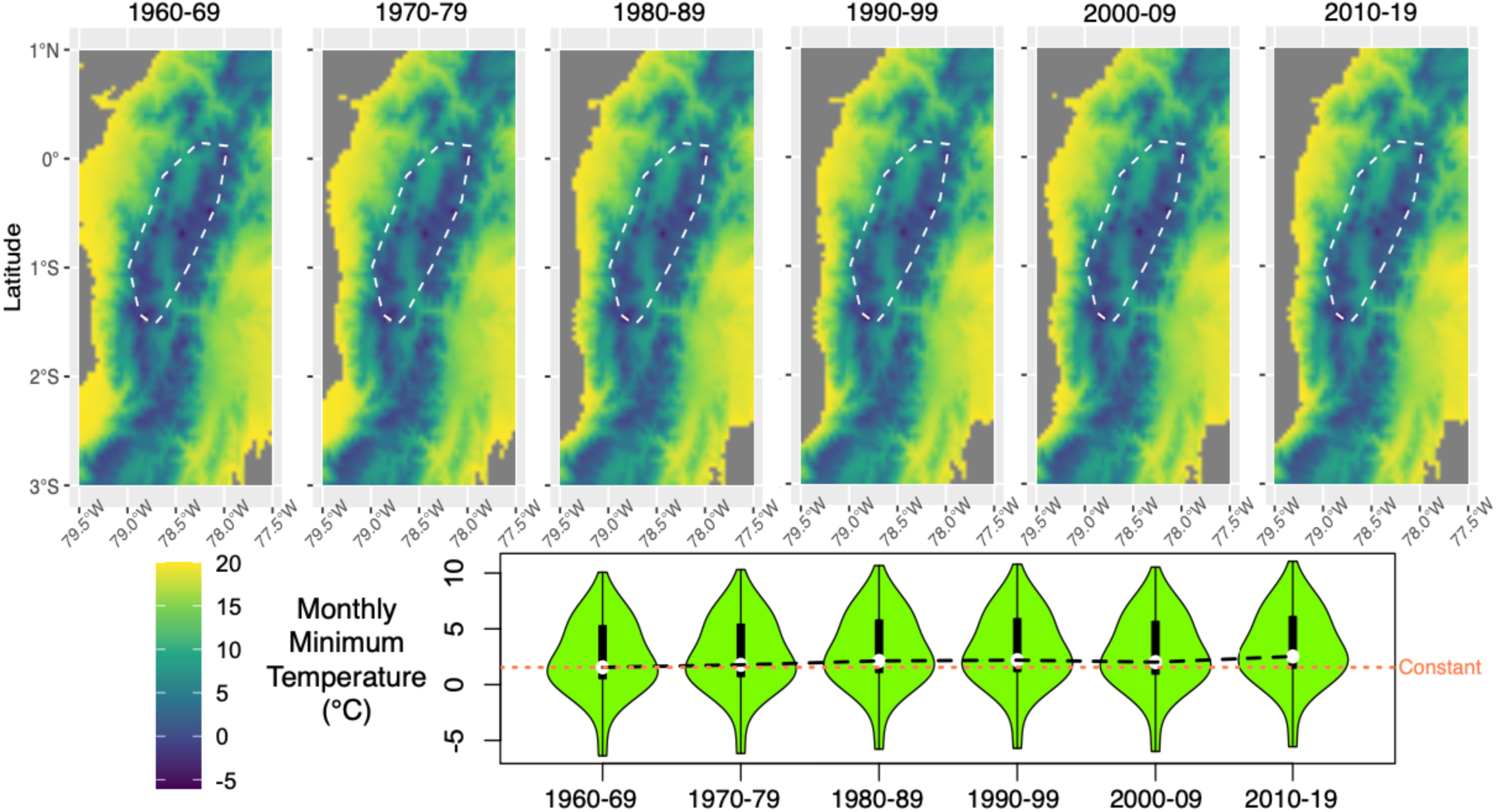
Annual mean of the monthly minimum temperature over decades. **Top**: The white dotted lines contour the *A. ignescens* known occurrence. **Bottom**: violin plot of monthly minimum temperature at all sites where the species was present. The median of the minimum temperature distribution increased over decades, deviating from the constant expectation line (dotted, in orange). The increase in minimum temperature was significant (*p* < 0.001, repeated measures ANOVA).

**Fig. 3.**
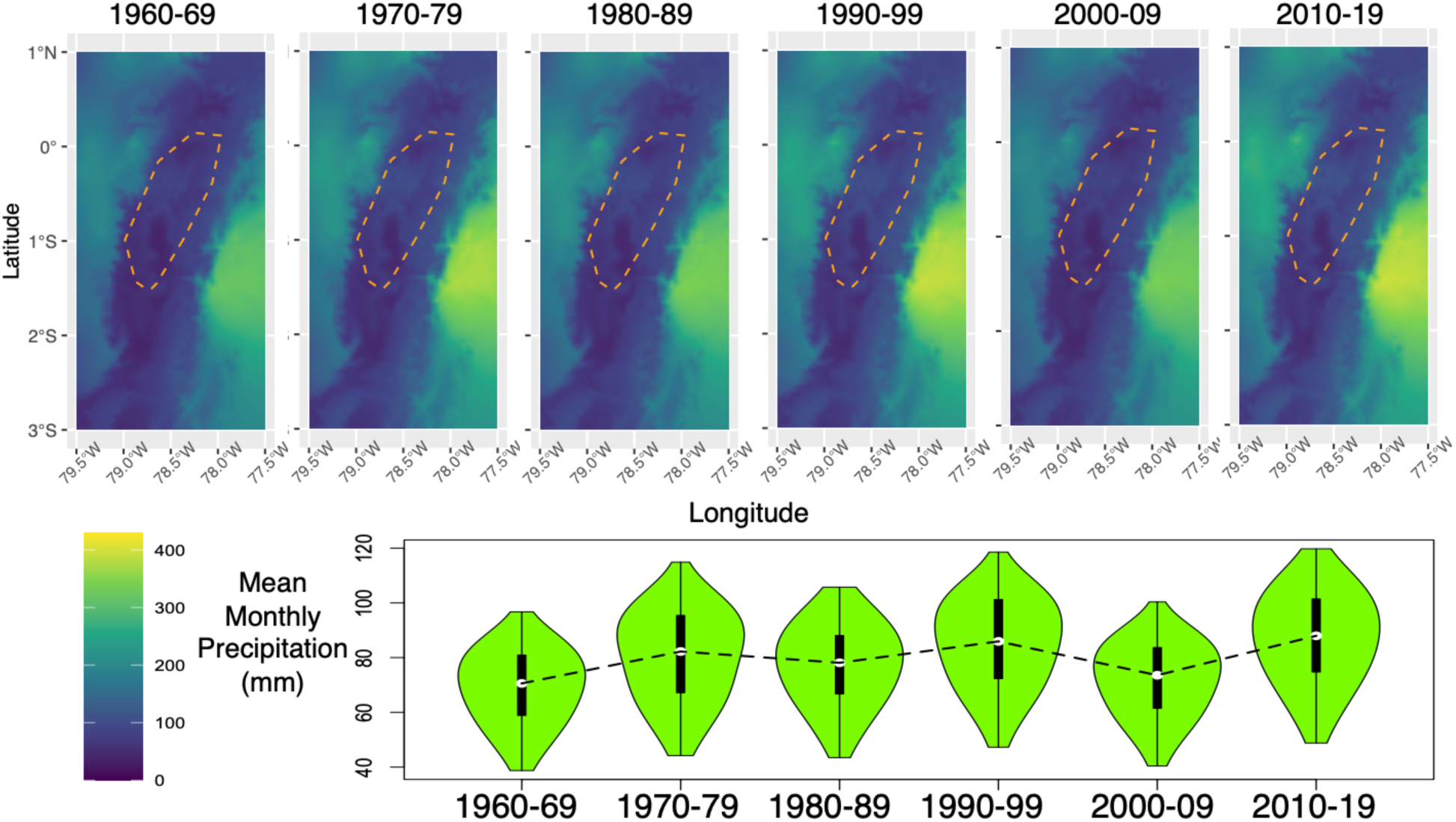
Annual mean of monthly precipitation (mm) over decades. **Top**: spatial distribution of precipitations. The orange contour outlines the *A. ignescens* known occurrence. **Bottom**: violin plots precipitation in sites where the species occurred. There was a significant change in precipitation over decades (*p* < 0.001, repeated measures ANOVA).

**Fig. 4.**
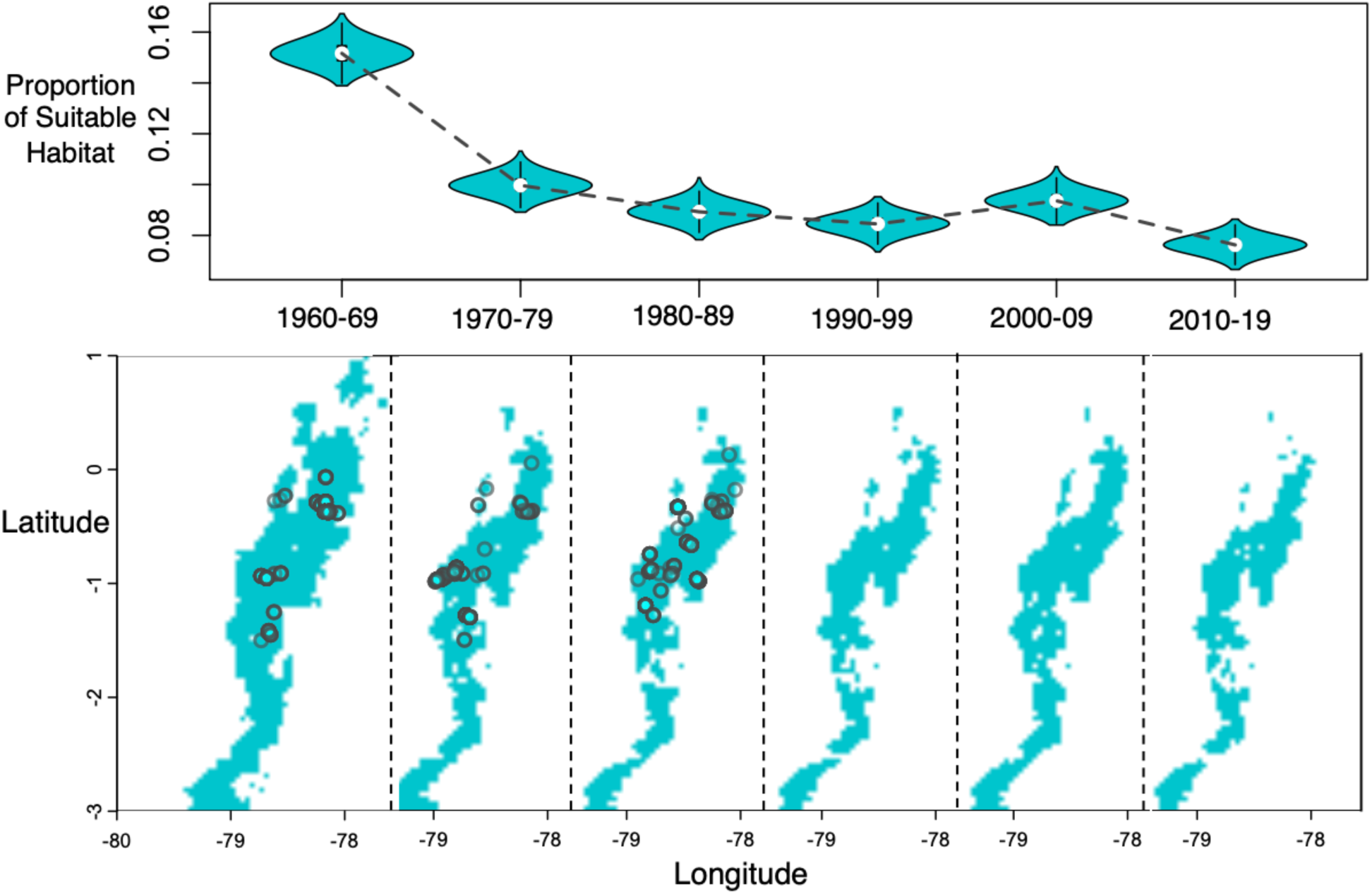
Predicted occurrence of *Atelopus ignescens* based on the threshold probability of presence. **Top**: violin plot of the bootstrap distribution of the proportion of suitable habitat, the proportion of area with expected presence based on the 1960-69 SDM, and climate data from each decade. **Bottom**: The turquoise area represents the area with predicted species presence, where the species distribution probability was greater than the estimated occurrence threshold of each decade. The reduction in the predicted area of occurrence is roughly 8190 km^2^ by year 2010-19. The black circles were historical sightings of *A. ignescens*.

## Discussion

We observed drastic shrinkage of the predicted distribution of *A. ignescens* during and after the rapid population decline in 1985 (Fig. 4). This phenomenon can be attributed to the changes in temperature (Fig. 2) and precipitation (Fig. 3) recorded in the Ecuadorian Andes after 1970 (Table S4). Therefore, climate change could have contributed strongly to the rampant decline of this species during the 1980s. There was a significant but subtle increase in male body size during *Bd* epizootics, which indicates that Bd infection could have affected the population before the historical extinction (Fig. 1 B, C). The excessive rare alleles in *A. ignescens* were consistent with a population bottleneck before the historical extinction, which was likely contributed by a reduction in habitat suitability due to climate change.

Temperature, precipitation, and their interactions explained over 72% of A. ignescens distribution during 1960-69 with 90% accuracy. This model effectively captures the subsequent climate changes (Fig. 2-3) and the impact on the decline in the predicted presence of *A. ignescens* (Fig. 4). While Bd is recognized as one of the main drivers of amphibian population decline (Lips 2008, Scheele 2019), some studies suggest a potential interplay between climate change and Bd. Certain environmental conditions may favor the fungus (Pounds 2006, Lips 2016, Ron 2003). Higher temperatures increase water vapor, which leads to greater cloud cover. This, in turn, could reduce nocturnal heat loss, resulting in nighttime warming and moderating daytime temperatures by blocking solar radiation. Consequently, the daily temperature range decreases as minimum temperatures rise, warm or dry conditions may stress amphibians, making them more susceptible to disease, while warmer years may directly benefit *Batrachochytrium* (Pounds et al. 2006).

One caveat is that the model prediction is the assumption of a constant climate-species relationship. The prediction accuracy declined over decades, reflecting distribution stochasticity and/or organismal response to climate change (Wang, 2024). There could have been plastic or evolutionary change in *A. ignescens* range in response to climate change since 1970. Future studies could examine whether such shifts were adaptive or maladaptive before the historical extinction.

Knowledge gap remains in the potential influence of seasonal climate (including dry periods), pathogen, additional latent factors, and their interactions on the decline of *A. ignescens*, as suggested by Ron et al. (2003), Merino-Viteri et al. (2005) and Menéndez and Graham (2013). Studies in other *Atelopus* species from Venezuela, identified an unusually dry year in 1988 when Bd-positive individuals were collected (Lampo et al. 2006). Despite this, the annual rainfall patterns did not show a clear relationship with the number of infected specimens detected.

Previous research on museum specimens revealed that most *A. ignescens* tested negative for Bd during the 1980s (Merino-Viteri 2001), and those that did test positive, had very low load (Manzano 2014). Our study suggests a slight influence of the Bd infection on the *A. ignescens* populations, as indicated by the larger male frogs observed during the 1980s compared to earlier periods. Smaller individuals seem to be more susceptible to chytridiomycosis, potentially due to the allometric scaling of metabolic rate and cutaneous ion loss rates, which makes younger and smaller frogs more vulnerable to osmoregulatory and metabolic disturbances (White et al. 2006, Wu et al. 2018). The frequency of sloughing observed in small frogs and those with significant Bd infections could have long-term effects on their ability to maintain homeostasis over time (Wu et al 2018). As a result, Bd epizootics might have exerted positive selection for larger body sizes during the 1980s. However, it is essential to acknowledge that other ecological and environmental factors could also play a role in selecting for larger body sizes and should be considered in future studies.

While massive die-offs occurred during the 1980s–1990s, not all Bd-infected species experienced population declines. Recent findings by Yánez et al. (2024) reported the presence of Bd in the only surviving population of *A. ignescens* in Angamarca. Remarkably, none of the infected individuals displayed visible signs of illness, suggesting a degree of resilience within this population despite infection. Bd susceptibility among Andean amphibian species varies widely, ranging from tolerance to high vulnerability (Catennazi et al. 2017; Guayasamin et al. 2014; Coloma & Duellman 2024). The interaction between the fungus Bd and Andean amphibians is highly complex and seems to vary significantly between species or populations. While some amphibian species or populations are severely affected by Bd, experiencing population declines or extinctions, others have shown the ability to survive and maintain stable populations. This variability underscores the adaptability and resilience of certain species, highlighting the need to consider species-specific and population-specific factors when studying the impacts of this widespread pathogen.

Consistently, we observed excessive rare alleles and marginal elevation of male body sizes during Bd epizootics. It is worth noting that Lampo et al. (2023) found that individuals of *A. cruciger* start to breed earlier in their lifecycle —at smaller body sizes— post-*Bd* epizootics. Unfortunately, we were unable to measure *A. ignescens* individuals after 1988 due to the lack of specimens in museum collections. Lampo’s hypothesis could potentially apply to *A. ignescens*, and further analysis should focus on the only currently known population of the Jambato Harlequin frog in Ecuador. Such a study could provide valuable insights into the role of Bd in the extinction of this species.

Climate change could have interacted with pathogens that led to the drastic decline of *A. ignescens*. Severe droughts have the potential to instigate chytridiomycosis outbreaks by amplifying the transmission rates of Bd or rendering the host vulnerability to the impacts of the pathogen (Daszak et al. 2003). Therefore, incorporating microclimate data from specific time points, such as 1988, within the projected distribution of the Jambato Harlequin frogs, would greatly benefit further research endeavors.

### Remaining knowledge gap & museum-omics

The remaining obstacles for Andean frog conservation are the *Bd* fungus and climate change. Understanding the response of the *Atelopus* frogs to these ongoing threats is the foundation for conservation. Genomic sequencing with museum specimens could unravel the population trend, potential climate adaptation, *Bd* immunity, and the genetic diversity of the *Atelopus* frogs. In the face of global warming, Bd fungus might be suppressed by suboptimal temperatures in certain geographic areas. However, the short generation time might allow the pathogens to rapidly adapt. The shift in male body size indicates the directional selection driven by *Bd* fungus epizootics. If there is enough genetic diversity for the *Atelopus* frogs to adapt to the pathogens, there is hope for *Atelopus* conservation. Museum curation harbors the opportunity to understand the genomics in both the pathogens and the hosts over space and time to inform the future of the resurrected frog species.

The examination of specimens of *A. ignescens* in biological collections has allowed us to gain insights into the historical rapid population decline. Synthesizing the limited information and museum specimens available is crucial for effective conservation strategies. By understanding the mechanisms underlying the population decline, we can develop optimal conservation strategies to protect this endangered species. Here, we show the significant relevance of museum specimens as a source of multidimensional information about the environment at specific times and places in the past. Museum data represent a significant tool for conservation (Nakahama 2021), offering valuable information essential for understanding and protecting endangered species such as *A. ignescens*.

## Acknowledgment

We thank Kevin Robert Chovanec (Kansas University) for accessing and measuring snout-vent lengths of 15 vouchered specimens. We thank Derek Eddo for formatting and proofreading part of the manuscript. This work is funded by NY SUTRA 960028-01 and SUNY RF 1183754-75023 to SW.

