## Supplement for "Museum insights for conservation: Unraveling the Extinction Factors in the Jambato Harlequin Frog, *Atelopus ignescens*"

**Supplementary Tables**

**Table S1 Individual IDs corresponding to GenBank accession ID.**

| ID | GenBank accession number | Voucher number | Gene name |
|---|---|---|---|
| **Indv 1** | **HM245262.1** | **KU 201106** | 16S ribosomal RNA gene (partial); tRNA-Leu gene (complete); NADH dehydrogenase subunit I gene (partial); tRNA-Ile gene (partial) |
| **Indv 2** | **DQ068444.1** | **KU 217433** | ADH dehydrogenase subunit 2 gene (partial); tRNA-Trp, tRNA-Ala, tRNA-Asx, tRNA-Cys, tRNA-Tyr genes (complete); cytochrome oxidase subunit I gene (partial) |
| **Indv 3** | **AY995950.1** | **KU 217433** | **cytochrome b gene** (partial) |
| **Indv 4** | **HM245261.1** | **WED 54159** | 16S ribosomal RNA gene (partial); tRNA-Leu gene (complete); NADH dehydrogenase subunit I gene (partial); tRNA-Ile gene (complete) |
| **Indv 5** | **HM245251.1** | **KU 201129** | 16S ribosomal RNA gene (partial); tRNA-Leu gene (complete); NADH dehydrogenase subunit I gene (partial); tRNA-Ile gene (partial) |
| **Indv 6** | **HM245254.1** | **KU 201125** | 16S ribosomal RNA gene (partial); tRNA-Leu gene (complete); NADH dehydrogenase subunit I gene (partial); tRNA-Ile |

| | | | |
|---|---|---|---|
| | | | gene (partial) |
| **Indv 7** | **HM245257.1** | **KU 201078** | 16S ribosomal RNA gene (partial); tRNA-Leu gene (complete); NADH dehydrogenase subunit I gene (partial); tRNA-Ile gene (partial) |
| **Indv 8** | **HM245264.1** | **KU 201132** | 16S ribosomal RNA gene (partial); tRNA-Leu gene (complete); NADH dehydrogenase subunit I gene (partial); tRNA-Ile gene (partial) |
| **Indv 9** | **HM245265.1** | **KU 201184** | 16S ribosomal RNA gene (partial); tRNA-Leu gene (complete); NADH dehydrogenase subunit I gene (partial); tRNA-Ile gene (partial) |
| **Indv 10** | **HM245266.1** | **KU 202200** | 16S ribosomal RNA gene (partial); tRNA-Leu gene (complete); NADH dehydrogenase subunit I gene (partial); tRNA-Ile gene (partial) |
| **Indv 11** | **HM245252.1** | **KU 201122** | 16S ribosomal RNA gene (partial); tRNA-Leu gene (complete); NADH dehydrogenase subunit I gene (partial); tRNA-Ile gene (partial) |
| **Indv 12** | **HM245253.1** | **KU 201124** | 16S ribosomal RNA gene (partial); tRNA-Leu gene (complete); NADH dehydrogenase subunit I-like, tRNA-Ile genes (partial) |
| **Indv 13** | **HM245255.1** | **KU 201076** | 16S ribosomal RNA gene (partial); tRNA-Leu gene |

| | | | |
|---|---|---|---|
| | | | (complete); NADH dehydrogenase subunit I gene (partial); tRNA-Ile gene (partial) |
| **Indv 14** | **HM245258.1** | **KU 201104** | 16S ribosomal RNA gene (partial); tRNA-Leu gene (complete); NADH dehydrogenase subunit I gene (partial); tRNA-Ile gene (partial) |
| **Indv 15** | **HM245262.1** | **KU 201106** | 16S ribosomal RNA gene (partial); tRNA-Leu gene (complete); NADH dehydrogenase subunit I gene (partial); tRNA-Ile gene (partial) |
| **Indv 16** | **HM245263.1** | **KU 201131** | 16S ribosomal RNA gene (partial); tRNA-Leu gene (complete); NADH dehydrogenase subunit I gene (partial); tRNA-Ile gene (partial) |

**Table S2** Stepwise model selection investigating the effects of sex, the time period before or during Bd infection (period), geographic index (geo), and their interactions (:) on SVLs of *A. ignescens*. The model (bolded) with the interaction of sex and periods explained the model the best (lowest AIC).

| **Predictors** | **AIC** |
|---|---|
| sex + period + geo + sex:period + sex:geo + period:geo + sex:period:geo | 507.30 |
| sex + period + geo + sex:period + sex:geo + period:geo | 505.11 |
| sex + period + geo + sex:period + sex:geo | 503.16 |
| sex + period + geo + sex:geo | 503.47 |

**Table S3** Stepwise model selection in species distribution model for 1960-1970

| Steps | Predictors | AIC |
|---|---|---|
| Start | Tmin + Tmax + Prec + Tmin:Tmax + Tmin:Prec + Tmax:Prec + Tmin:Tmax:Prec | 249.38 |
| Step1 | Tmin + Tmax + Prec + Tmin:Tmax + Tmin:Prec + Tmax:Prec | 248.65 |
| Step 2 | Tmin + Tmax + Prec + Tmin:Tmax + Tmin:Prec | 247.17 |

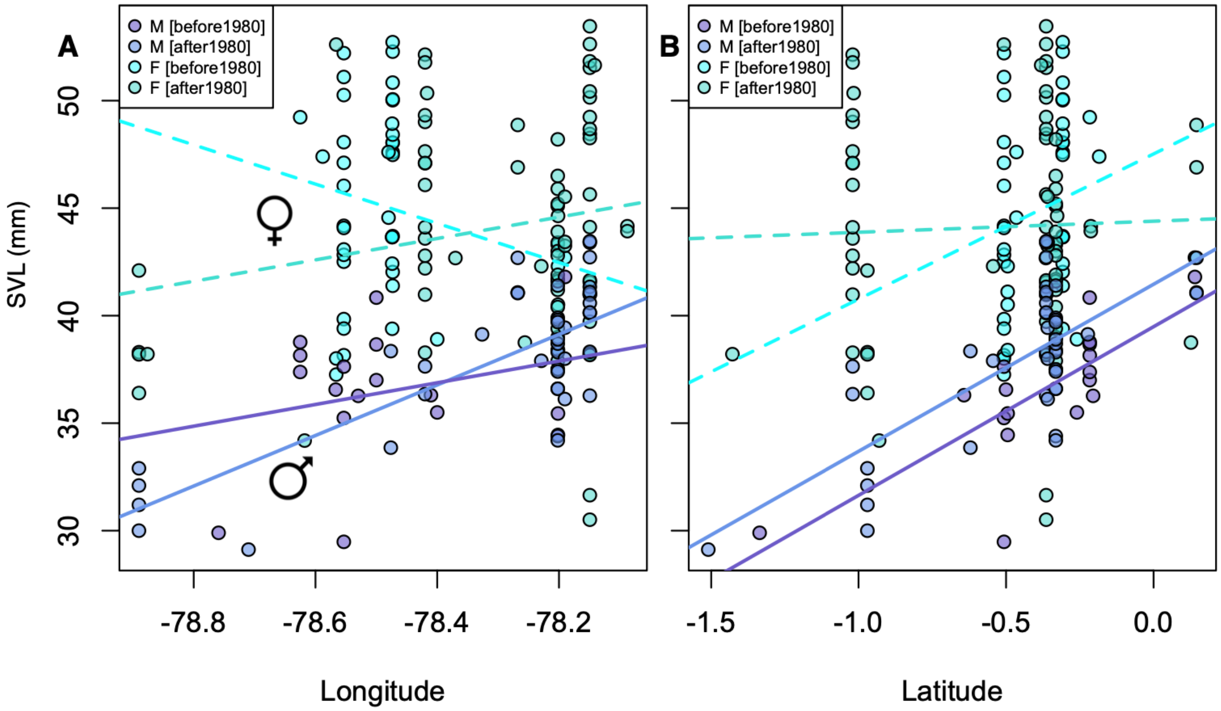


**Fig. S1 The effects of sex, geography, and periods (before vs after 1980) on SVL (mm). A-B,** Females have much higher SVL than males. The SVL forms geographic clines in both time periods when the individuals in the higher longitude (A) and higher latitude **(B)** are larger than individuals in the southwest. Females did not exhibit SVL clines over geographic space.

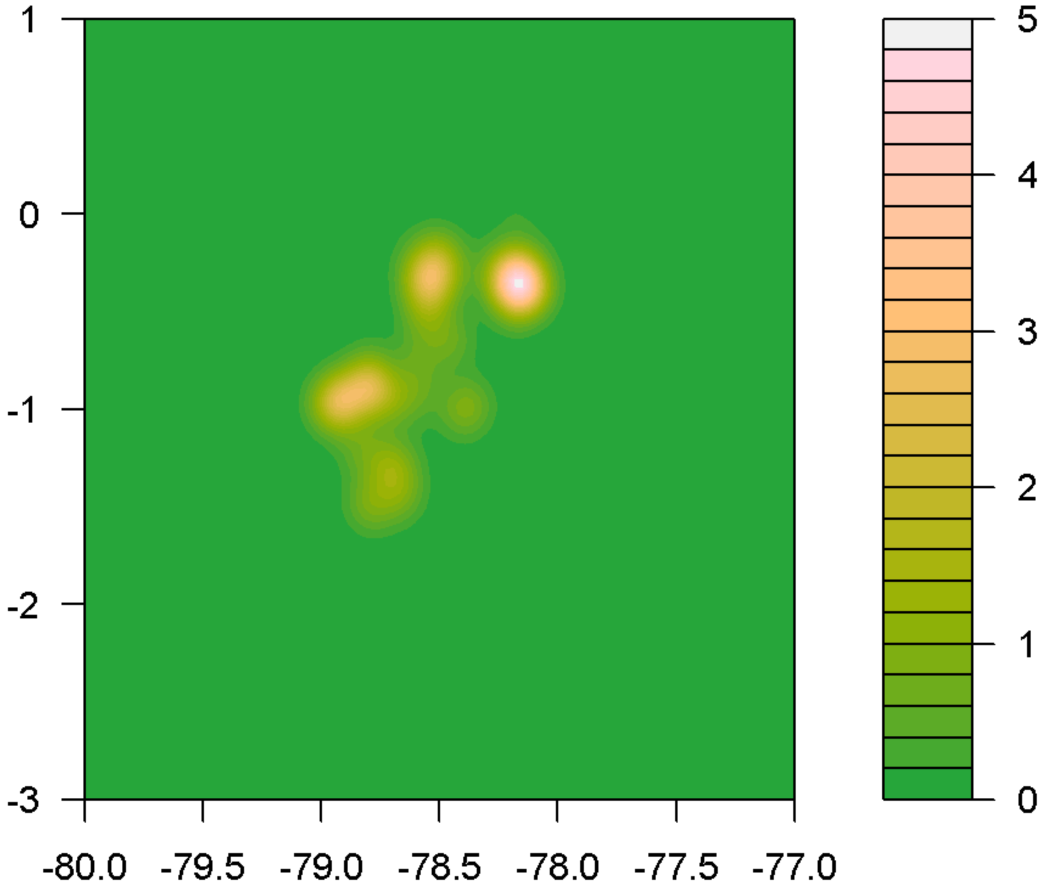


**Fig. S2**. Contour plot of *A. ignescens* density based on presence data of all time (N =1122). The density kernel is approximated with bivariate normal distribution. The x-axis and y-axis are respectively the longitude and latitude.

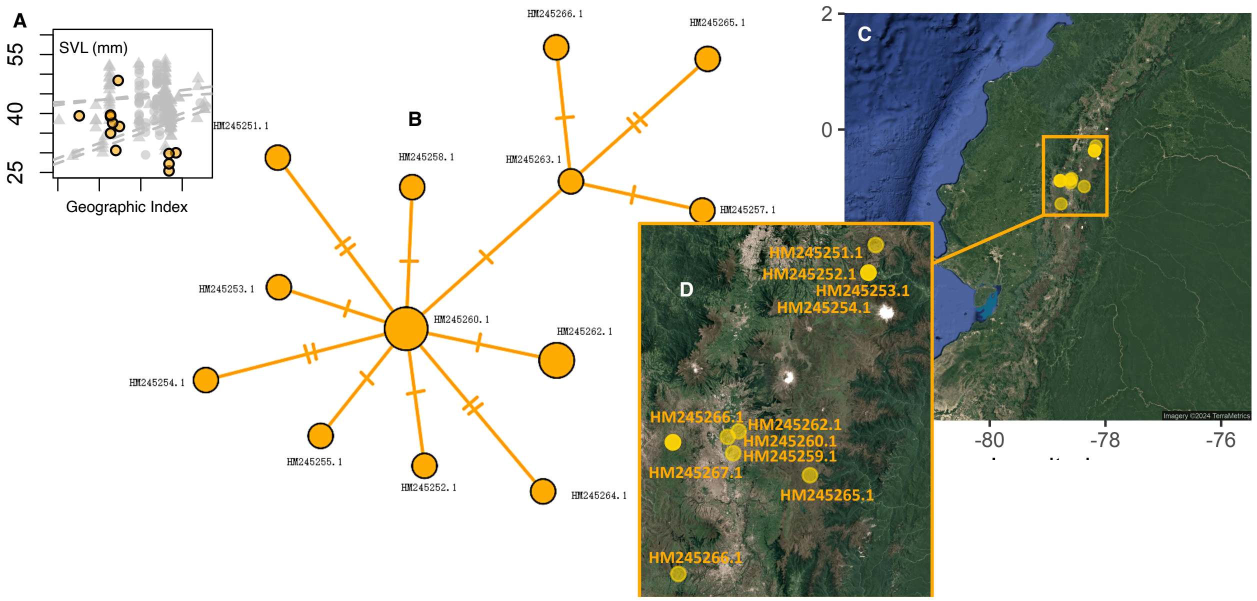


**Fig. S3 Geographic location corresponding to the nodes in the haplotype network (Fig. 1 D). A**, SVL, and geographic index among the genotyped individuals relative to the individuals analyzed in **Fig. 1B** colored in grey, haplotype network with Genbank ID (Table S1). **C**, a map showing the geographic location where the mtDNA samples were from. The haplotype groups do not represent spatial groups.

**Table 4 Repeated Measure ANOVA** of climate change among time period in sites where *A. ignescens* occurred.

| | $F_{(5, 5605)}$ | *p-value* |
|---|---|---|
| **Tmax** | 80733.1966 | <0.00001 |
| **Tmin** | 16331.16 | <0.00001 |
| **Tprec** | 15650.49 | <0.00001 |
